# Effect of Positive Allosteric Modulation and Orthosteric Agonism of Dopamine D2 Receptors on Respiration in Mouse Models of Rett Syndrome

**DOI:** 10.1101/2022.04.13.488220

**Authors:** Sebastian N. Maletz, Brandon T. Reid, David M. Baekey, Jessica R. Whitaker-Fornek, Jordan T. Bateman, John M. Bissonnette, Erica S. Levitt

**Affiliations:** Department of Pharmacology and Therapeutics, University of Florida, Gainesville, FL 32610; Breathing Research and Therapeutics Center, University of Florida, Gainesville, FL 32610; Oregon Health and Sciences University, Portland, OR 97239

**Keywords:** Dopamine D2 receptors, Rett Syndrome, respiration, electrophysiology

## Abstract

Rett syndrome (RTT) is an autism spectrum disorder caused by loss-of-function mutations in the methyl-CPG-binding protein 2 (Mecp2) gene. Frequent apneas and irregular breathing are prevalent in RTT, and also occur in rodent models of the disorder, including Mecp2^Bird^ and Mecp2^R168X^ mice. Sarizotan, a serotonin 5-HT1a and dopamine D2-like receptor agonist, reduces the incidence of apneas and irregular breathing in mouse models of RTT (Abdala et al., 2014). Targeting the 5HT1a receptor alone also improves respiration in RTT mice (Levitt et al., 2013). However, the contribution of D2 receptors in correcting these respiratory disturbances remains untested. PAOPA, a dopamine D2 receptor positive allosteric modulator, and quinpirole, a dopamine D2 receptor orthosteric agonist, were used in conjunction with whole-body plethysmography to evaluate whether activation of D2 receptors is sufficient to improve breathing disturbances in female heterozygous Mecp2^Bird/+^ and Mecp2^R168X/+^ mice. PAOPA did not significantly change apnea incidence or irregularity score in RTT mice. PAOPA also had no effect on the ventilatory response to hypercapnia (7% CO_2_). In contrast, quinpirole reduced apnea incidence and irregularity scores and improved the hypercapnic ventilatory response in Mecp2^R168X/+^ and Mecp2^Bird/+^ mice, while also reducing respiratory rate. These results suggest that D2 receptors do contribute to the positive effects of sarizotan in the correction of respiratory abnormalities in Rett syndrome. However, positive allosteric modulation of the D2 receptor alone is not sufficient to evoke these effects.

## Introduction

Rett syndrome (RTT) is an X-linked neurodevelopmental disorder afflicting 1:10,000-1:15,000 females that presents with neurological cognitive and motor deficits, anxiety, seizures and breathing dysfunction (Smeets *et al*., 2012). RTT is caused by loss-of-function mutations in the gene that encodes the transcription factor methyl-CpG-binding protein 2 (Mecp2) (Amir *et al*., 1999; Guy *et al*., 2011). Respiratory disturbances, including apneas, hyperventilation, an irregular breath cycle, and excessive air swallowing occur in nearly all patients with RTT (Ramirez *et al*., 2013; Mackay *et al*., 2017; Tarquinio *et al*., 2018). These respiratory abnormalities, in the form of apnea and irregular breathing, are also observed in rodent models of RTT, including Mecp2^Bird/+^ and Mecp2^R168X/+^ mice (Levitt *et al*., 2013*a*; Abdala *et al*., 2014). The mixed serotonin 5HT1a and dopamine D2 receptor agonist sarizotan corrects these breathing disturbances in Mecp2^Bird/+^ and Mecp2^R168X/+^ mice (Abdala *et al*., 2014). Selective 5HT1A receptor agonists also correct breathing irregularity in Mecp2^Bird/+^ mice (Levitt *et al*., 2013*a*). The contribution of D2 receptors to the positive effects of sarizotan remains untested.

Apneas that occur in mouse models of RTT are associated with augmented discharge from expiratory nerves (Abdala *et al*., 2010). Dopamine D2 receptors suppress the activity of expiratory neurons in the caudal ventral respiratory group (Lalley, 2009), and are thus potential targets to correct the apneas and breathing irregularity observed in RTT. D2 receptor agonists are currently used clinically as therapeutics for Parkinson’s disease (BROOKS, 2000). Several compounds that act allosterically at the dopamine D2 receptor are also under development (Wood *et al*., 2016; Wold *et al*., 2019; Fasciani *et al*., 2020). The objective of the present study was to determine the impacts of compounds that enhance D2 receptor activity on respiration in mouse models of RTT. Respiration was evaluated in normal air and also during hypercapnic challenge, since Mecp2 deficient mice have an impaired hypercapnic ventilatory response (Zhang *et al*., 2011; Toward *et al*., 2013; Bissonnette *et al*., 2014; Garg *et al*., 2015).

In this study, we used two different compounds selectively targeting the dopamine D2 receptor. PAOPA, (3R)-2-Oxo-3-[[(2S)-2-Pyrrolidinylcarbonyl]amino]-1-pyrrolidineacetamide, is a potent and specific positive allosteric modulator of the dopamine D2 receptor, and is being tested in preclinical models of tardive dyskinesia, schizophrenia and Parkinson’s disease (Marcotte *et al*., 1998; Sharma *et al*., 2003; Verma *et al*., 2005; Beyaert *et al*., 2013; Tan *et al*., 2013; Daya *et al*., 2018). The orthosteric dopamine D2 receptor agonist quinpirole was used for further investigation into effects of direct dopamine D2 receptor activation.

## Methods

### Ethical Approval

All experiments were approved by the Institutional Animal Care and Use Committee at the University of Florida and were in agreement with the National Institutes of Health “Guide for the Care and Use of Laboratory Animals.” Mice were bred and maintained at the University of Florida animal facility. Mice were group-housed in standard sized plastic cages and kept on a 12 h light–dark cycle, with water and food available *ad libitum*.

### Animals

Female heterozygous MeCP2 mutation knock-in mice, B6.129P2(C)-*Mecp2*^*tm1*.*1Bird*^ (stock no. 003890, Jackson Labs, Bar Harbor, ME) and B6J;129S6.*Mecp2*^*R168X*^ (stock no. 024990, Jackson Labs, Bar Harbor, ME), were purchased from Jackson Labs and backcrossed with male wild-type C57BL/6 mice for one generation (Mecp2^R168X^) or three generations (Mecp2^Bird^) to establish the colony, then maintained by crossing with wild-type males from the colony. Female heterozygous Mecp2-deficient Mecp2^Bird/+^ mice (ages 6-10 months) and Mecp2^R168X/+^ mice (ages 5-10 months) were used for plethysmography studies. Wild-type C57BL6/J mice (male and female, 2-4 months old) were used for brain slice electrophysiology experiments.

### Drugs

PAOPA ((3R)-2-Oxo-3-[[(2S)-2-Pyrrolidinylcarbonyl]amino]-1-pyrrolidineacetamide) was acquired from Tocris (Minneapolis, MN). (-)Quinpirole HCl and dopamine HCl were acquired from Sigma-Aldrich (St. Louis, MO). PAOPA and quinpirole for plethysmography studies were diluted in sterile saline. Drugs for brain slice experiments were dissolved in ultrapure water and diluted in ACSF at the indicated concentrations.

### Plethysmography Studies

Whole-body plethysmography (vivoFlow, SCIREQ Inc, Montreal, QC, Canada) was used to measure respiratory activity of awake mice. Mice were acclimated to the plethysmography chambers for three hours on three consecutive days prior to the first experimental day. On experimental days, mice were acclimated to the chambers for 15 minutes, then given saline (10 µl/g, i.p.), PAOPA (1 or 10 mg/kg in saline, i.p.), or quinpirole (1 mg/kg in saline, i.p.). For PAOPA experiments, each mouse was treated with saline, 1 mg/kg PAOPA, and 10 mg/kg PAOPA in a pseudo-random order across three experimental days, with a minimum interval of 3 days between doses. For quinpirole experiments, animals were treated with saline and then quinpirole one day later. Following drug or saline injection, mice were placed in amber-colored Plexiglas chambers which were ventilated with a constant flow (0.5 L/min) of compressed room air (21% O_2_, balance N_2_) for 60 minutes, followed by a 10-minute challenge with hypercapnic air (7% CO_2_, 21% O_2_, balance N_2_). Body surface temperature was measured using an infrared thermometer directed at the mid-sternal surface of the chest.

Data were analyzed for the period from 30-60 minutes post-injection of drug or saline in standard air (6,000-8,000 breaths), and the 10-minute period in hypercapnia. Respiratory frequency (f), inspiratory duration (T_i_) and expiratory duration (T_e_) of individual breaths were calculated using IOX2 software (SCIREQ Inc, Montreal, QC, Canada) and exported into Microsoft Excel. Events were rejected if the peak-to-peak rate failed to reach the event validation threshold of 1.0 ml/s or if the ratio of inspiratory volume to expiratory volume was below 70%. An apnea was defined as a breath with expiratory time greater than double the average total respiratory cycle time for that mouse (T_e_ ≥ 2 x average T_tot_). Irregularity score was defined as the difference in duration between two adjacent breaths divided by the duration of the first breath [(Ttot_n_-Ttot_n+1_)/Ttot_n+1_] and is reported as variance. Coefficient of variation (CoV) for various parameters was calculated using [(standard deviation/mean)*100].

### Brain Slice Electrophysiology

Mice were anesthetized with isoflurane and decapitated. The brain was removed and mounted in a vibratome chamber (Leica VT 1200S, Leica Biosystems, Buffalo Grove, IL, USA). Horizontal slices (230 µm) containing substantia nigra pars compacta were cut in artificial cerebrospinal fluid (ACSF) that contained the following (in mM): 126 NaCl, 2.5 KCl, 1.2 MgCl_2_, 2.4 CaCl_2_, 1.2 NaH_2_PO_4_, 11 D-glucose and 21.4 NaHCO_3_ (equilibrated with 95% O2/5% CO2). Slices were stored at 32°C in glass vials with equilibrated ACSF. MK801 (10 µM) was added to the cutting solution and for the initial incubation of slices in storage (at least 30 min) to block NMDA receptor-mediated excitotoxicity. Following incubation, the slices were transferred to a recording chamber that was perfused with equilibrated ACSF warmed to 34°C (Warner Instruments) and flowing at a rate of 1.5-3 ml/min.

Whole-cell recordings from substantia nigra pars compacta neurons were performed with a Multiclamp 700B amplifier (Molecular Devices, Sunnyvale, CA) in voltage-clamp mode (Vhold = -60mV) mode. Dopamine neurons within the substantia nigra pars compacta were identified based on location relative to the medial terminal nucleus of the accessory optic tract, large size, presence of Ih current and dopamine D2-receptor mediated GIRK current. Recording pipettes (1.5 – 3 MΩ) were filled with internal solution that contained (in mM): 115 potassium methanesulfonate, 20 NaCl, 1.5 MgCl_2_, 5 HEPES(K), 2 BAPTA, 1-2 Mg-ATP, 0.2 Na-GTP, adjusted to pH 7.35 and 275-285 mOsM. Data were low-pass filtered at 5 kHz and collected at 10 kHz with pClamp10.7 and at 400 Hz with PowerLab (Lab Chart version 5.4.2; AD Instruments, Colorado Springs, CO). Drugs (including quinpirole and PAOPA) were applied by bath perfusion at the indicated concentrations.

### Statistical Analysis

Statistical analyses were performed in GraphPad Prism 8. Data are reported as means ± SEM, with individual data points also shown whenever possible. Data with n > 8 were tested for normality with D’Agostino & Pearson tests. For non-normally distributed data, non-parametric tests were used: Wilcoxon test for comparisons between two groups of paired data, Friedman test with Dunn’s multiple comparisons test for comparisons between three or more groups of repeated measures data. For normally distributed data, parametric tests were used: two-tailed t-tests for comparisons between two groups, one-way ANOVA followed by Tukey’s post-hoc test or Holm-Sidak post hoc test for comparisons between three or more groups. Paired or repeated measures comparisons were made when appropriate, as indicated in Results.

## Results

### Validation of PAOPA as a Dopamine D2 Receptor Positive Allosteric Modulator

Activation of dopamine D2 receptors on midbrain dopamine neurons causes a robust GIRK-mediated outward current (Beckstead *et al*., 2004). We used this known D2-receptor mediated effect to validate the dopamine D2 receptor positive allosteric modulator PAOPA. Whole-cell voltage-clamp recordings were made from substantia nigra pars compacta (SNpc) dopamine neurons contained in brain slices from wild-type mice. Dopamine (10 µM) caused an outward current that was 66 ± 9 pA (n = 17; Figure 1A). Repeated application of dopamine (10 µM) produced outward currents of similar amplitude (98 ± 4 % of control (p = 0.547 Wilcoxon test, n = 8; Figure 1B)). PAOPA (30 µM) did not cause a current different from baseline (p = 0.189 paired t-test, n = 9), but enhanced the dopamine-mediated current to 157 ± 18 % of control (Figure 1A,C, p = 0.016 Wilcoxon test, n = 7), consistent with the expected properties of an allosteric agonist (Verma *et al*., 2005; Fasciani *et al*., 2020).

**Figure 1.**
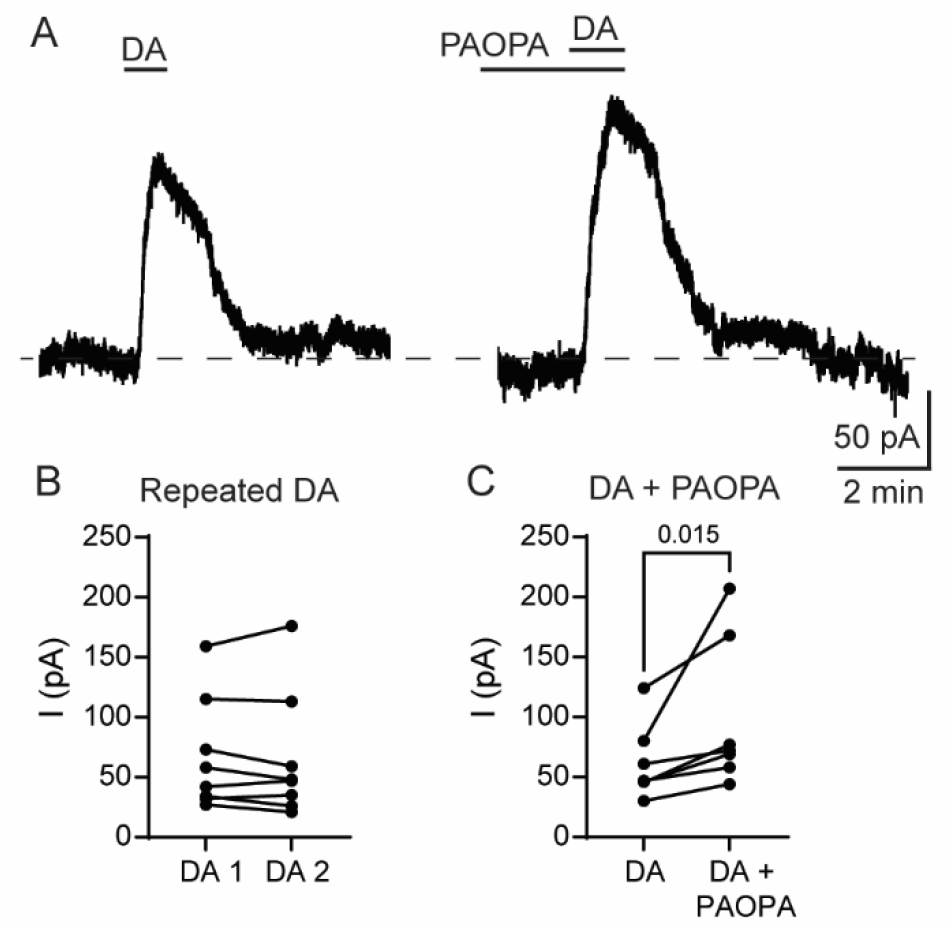
D2 allosteric modulator PAOPA enhances dopamine (DA)-mediated currents. A, Whole-cell voltage-clamp recording from midbrain dopamine neuron in brain slice from wild-type mouse. DA (10 µM) was applied before and during PAOPA (30 µM). B, Amplitude of current produced by repeated DA (10 µM). C, Amplitude of current produced by DA (10 µM) and DA (10 µM) + PAOPA (30 µM). p value from Wilcoxon test.

### Baseline Respiratory Patterns in Mecp2-deficient Mice

We used whole-body plethysmography to measure respiration in female heterozygous Mecp2^Bird/+^ and Mecp2^R168X/+^ mice that were 5-10 months old (n = 36 mice). Characteristic periodic breathing, marked by a waxing and waning pattern, was observed in both strains of Mecp2-deficient mice (Figure 2). Both strains of mice displayed apneas, defined as expiratory duration that was longer than two respiratory cycles (Table 1). The number of apneas in each strain was similar to previous reports (Levitt *et al*., 2013*b*; Abdala *et al*., 2014; Bissonnette *et al*., 2014). The irregularity in breathing pattern in MeCP2 mutant mice can also be analyzed on a breath-to-breath basis using irregularity score. Irregularity scores (Table 1) in Mecp2^R168X/+^ mice were similar to previously reported, while irregularity scores in Mecp2^Bird/+^ mice were about two-thirds of the previously reported value (Abdala *et al*., 2014). There was no difference in baseline apnea number (p = 0.1496 unpaired t-test), irregularity score (p = 0.5250 unpaired t-test) or average apnea length (p = 0.608 unpaired t-test) between the Mecp2^Bird/+^ or Mecp2^R168X/+^ mice that we tested (Table 1). Mecp2^Bird/+^ mice did have a higher respiratory frequency than Mecp2^R168X/+^ mice (p = 0.0099 unpaired t-test; Table 1).

**Figure 2.**
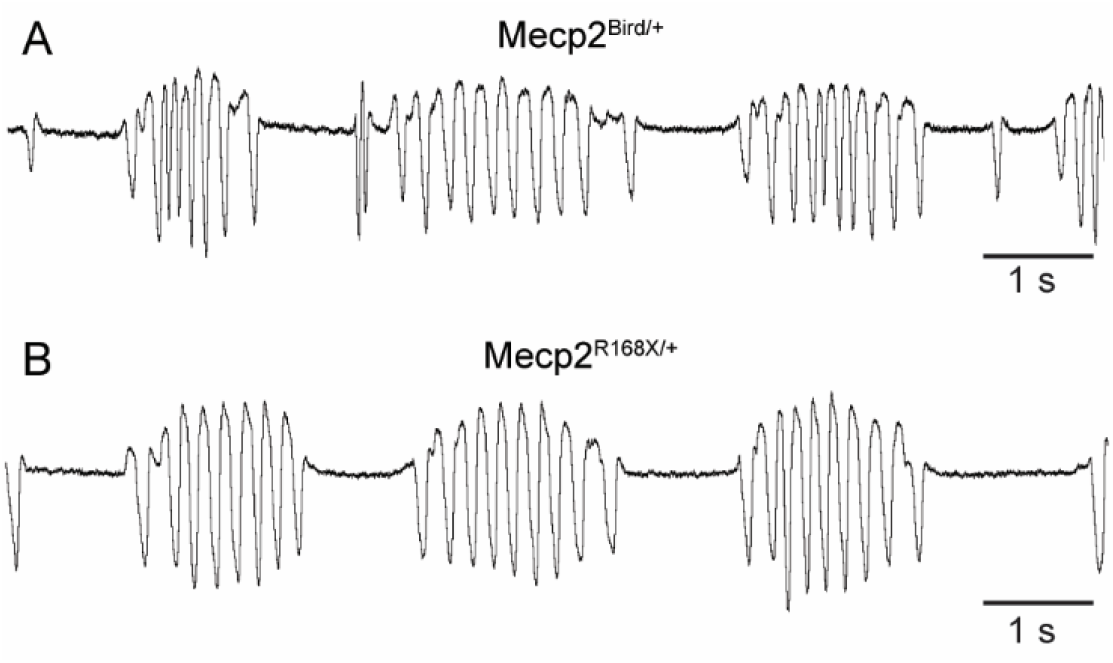
Representative whole body plethysmography traces from Mecp2^Bird/+^ (A) and Mecp2^R168X/+^ (B) mice showing apneas and periodic breathing.

**Table 1:** Baseline Respiratory Patterns in Mouse Models of Rett Syndrome.

| Strain | Frequency<br>(bpm) | Irregularity<br>(variance) | Apneas/hour | Apnea<br>Duration (ms) |
| --- | --- | --- | --- | --- |
| Mecp2<sup>Bird/+</sup> (n=18) | $382.8 \pm 48.8$ | $0.273 \pm 0.138$ | $230.2 \pm 198$ | $548.7 \pm 137$ |
| Mecp2<sup>R168X/+</sup> (n=18) | $333.9 \pm 58.3^*$ | $0.247 \pm 0.107$ | $147.0 \pm 134$ | $572.5 \pm 139$ |

We used linear regression to investigate the relationship between body weight (g) and apnea incidence or irregularity score. There was a significant negative correlation between body weight and apnea incidence in both strains of mice (Mecp2^Bird/+^ p<0.0001 and Mecp2^R168X/+^ p = 0.0135), and a significant negative correlation between body weight and irregularity score in both strains of mice ((Mecp2^Bird/+^ p < 0.0001 and Mecp2^R168X/+^ p = 0.0001), demonstrating that heavier mice had fewer apneas and less irregular breathing. There was no difference in mean body weight between the two strains of mice we tested (Mecp2^Bird/+^ 41.8 ± 3.3 g, Mecp2^R168X/+^ 45.8 ± 2.5 g, p = 0.3344 unpaired t-test). Thus, the irregular breathing pattern is unlikely due to fat accumulation and/or obesity in the MeCP2 mutant mice. In contrast, age was not significantly correlated with apnea incidence (Mecp2^Bird/+^ p = 0.1401 and Mecp2^R168X/+^ p = 0.0825) or irregularity score (Mecp2^Bird/+^ p = 0.1217 and Mecp2^R168X/+^ p = 0.0556) in either strain.

### Effects of Dopamine D2 Receptor Allosteric Modulator PAOPA on Respiration

We tested the ability of the dopamine D2 receptor positive allosteric modulator PAOPA (1 mg/kg and 10 mg/kg) to correct apneas and irregular breathing in both Mecp2^Bird/+^ (n = 8) and Mecp2^R168X/+^ (n = 10) mice. These doses of PAOPA have been previously reported to have behavioral effects lasting at least an hour following systemic administration in preclinical tests of sensorimotor deficits (Beyaert *et al*., 2013; Tan *et al*., 2013). Data were analyzed for the period 30-60 minutes following drug or vehicle injection. PAOPA (1 mg/kg and 10 mg/kg) did not significantly change the number of apneas or the irregularity score in either Mecp2^Bird/+^ or Mecp2^R168X/+^ mice (Figure 3). Further, there were no differences in average apnea duration between saline, 1 mg/kg and 10 mg/kg of PAOPA treatment (Mecp2^Bird/+^: 505 ± 44 ms (saline), 497 ± 52 ms (1 mg/kg), 489 ± 44 ms (10 mg/kg), p = 0.879 by one-way ANOVA (p > 0.05 for all multiple comparisons by Tukey post-test)); Mecp2^R168X/+^, 536 ± 47 ms (saline), 520 ± 49 ms (1 mg/kg), 550 ± 49 ms (10 mg/kg), p = 0.499 by one-way ANOVA (p > 0.05 for all multiple comparisons by Tukey post-test)). Respiratory frequency, inspiratory duration, expiratory duration, and tidal volume were all unchanged by PAOPA treatment (Figure 4).

**Figure 3.**
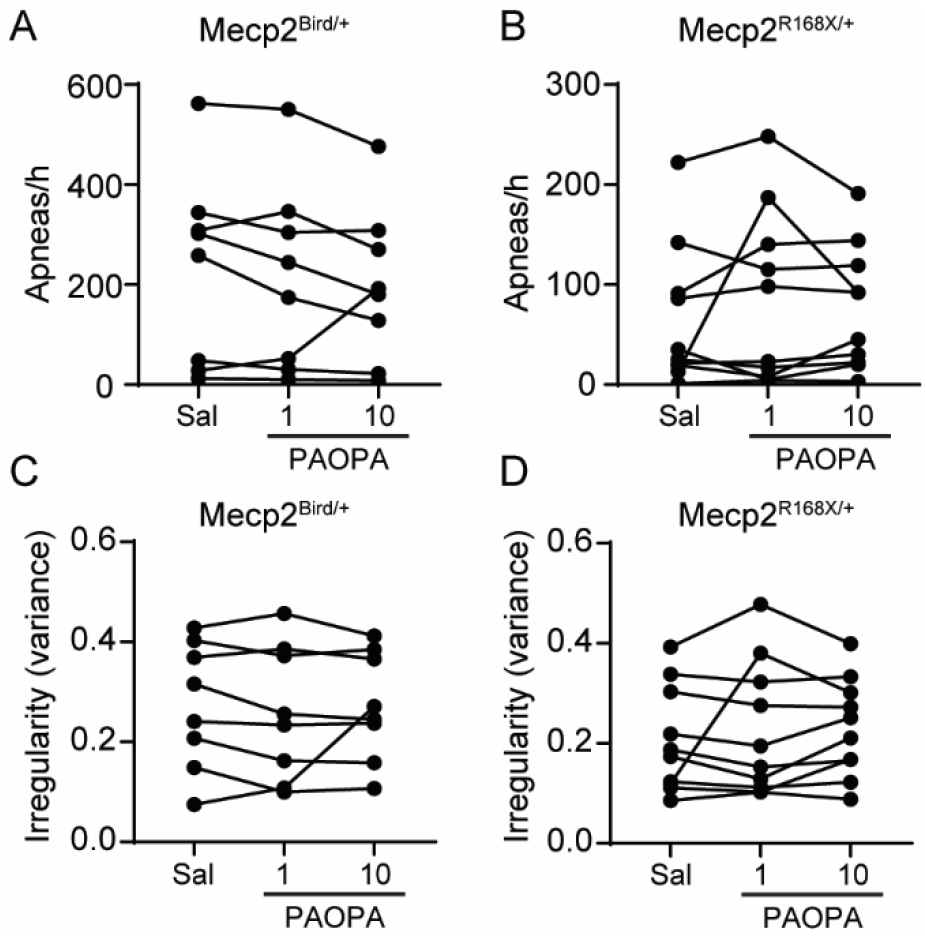
Apnea and irregularity scores were unchanged by PAOPA. Whole-body plethysmography data from Mecp2^Bird/+^ or Mecp2^R168X/+^ mice were analyzed 30 – 60 min after saline or PAOPA (1 or 10 mg/kg). Data points and connecting lines represent means from individual animals. For all parameters no significant differences were detected (p > 0.05 by RM one-way ANOVA and Tukey multiple comparisons post-test). RM ANOVA p-values were as follows: Apnea/h: Mecp2^Bird/+^ p = 0.377, Mecp2^R168X/+^ p = 0.397; Irregularity: Mecp2^Bird/+^ p = 0.684, Mecp2^R168X/+^ p = 0.455.

**Figure 4.**
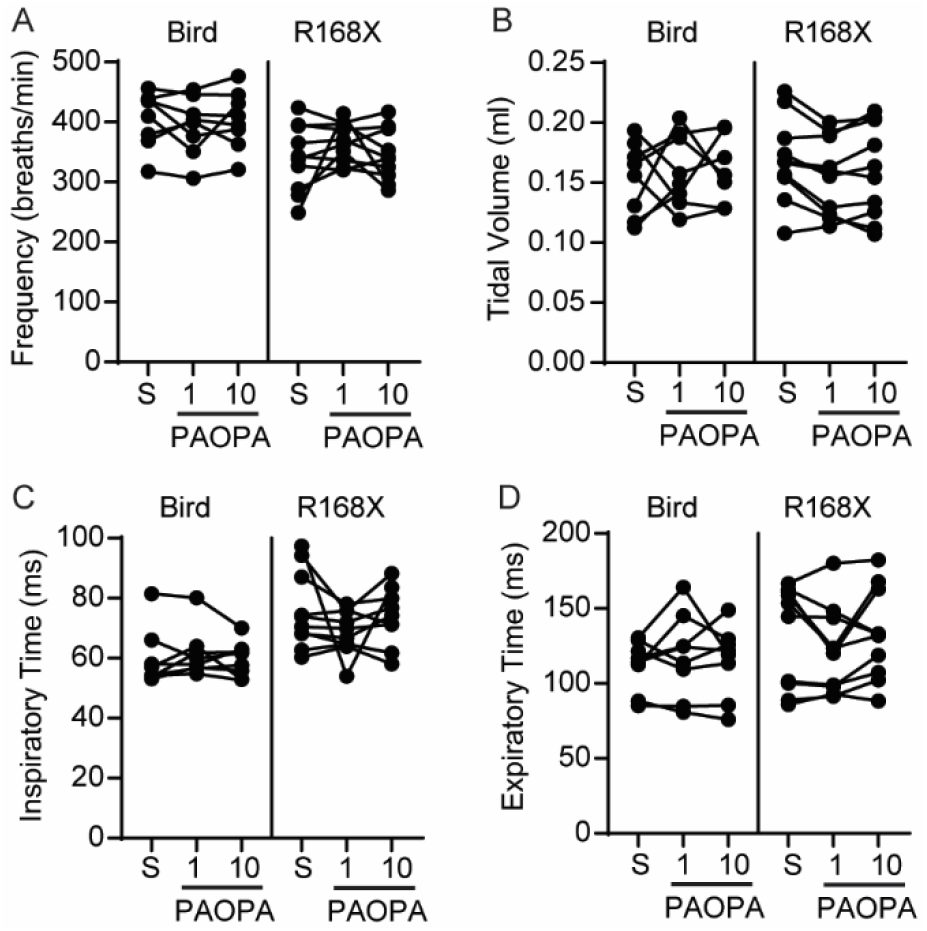
Whole-body plethysmography data from Mecp2^Bird/+^ or Mecp2^R168X/+^ mice. Respiratory parameters were measured 30-60 minutes following injection of saline or PAOPA (1 or 10 mg/kg). Data points and connecting lines represent means from individual animals. For all parameters no significant differences were detected (p > 0.05 by RM one-way ANOVA and Tukey post-test).

### Effects of Dopamine D2 Receptor Agonist Quinpirole on Respiration

Since we did not observe effects with the positive allosteric D2 modulator PAOPA, we tested the orthosteric D2 agonist quinpirole to determine if direct activation of the dopamine D2 receptor would improve apneas and irregular breathing. Mice (Mecp2^Bird/+^ (n = 10) and (Mecp2^R168X/+^ (n = 8)) were injected with saline or quinpirole (1 mg/kg). Data were analyzed 30-60 minutes following injection. Apnea incidence was significantly reduced following quinpirole treatment in both Mecp2^Bird/+^ and Mecp2^R168X/+^ mice relative to saline (Figure 5A). The average length of apneas that did occur was not changed by quinpirole treatment (Mecp2^Bird/+^ 584 ± 45 ms (saline) vs 775 ± 63 ms (quinpirole), p = 0.0612 by paired t-test; Mecp2^R168X/+^ 618 ± 42 ms (saline) to 746 ± 81 ms (quinpirole), p = 0.1773, paired t-test). Quinpirole administration also significantly reduced irregularity scores in both Mecp2^Bird/+^ and Mecp2^R168X/+^ mice (Figure 5B). However, quinpirole also had effects on baseline breathing (Figure 6) and body temperature (saline = 36.3 ± 0.7 ^O^F; quinpirole = 34.5 ± -.12 ^O^F; p = 0.0004 paired t-test, n = 5). Breathing frequency and tidal volume was reduced in quinpirole-treated mice (Figure 6 A,B). The reduction in frequency observed in Mecp2^Bird/+^ mice was driven by an increase in both inspiratory and expiratory time (Figure 6 C,D).

**Figure 5.**
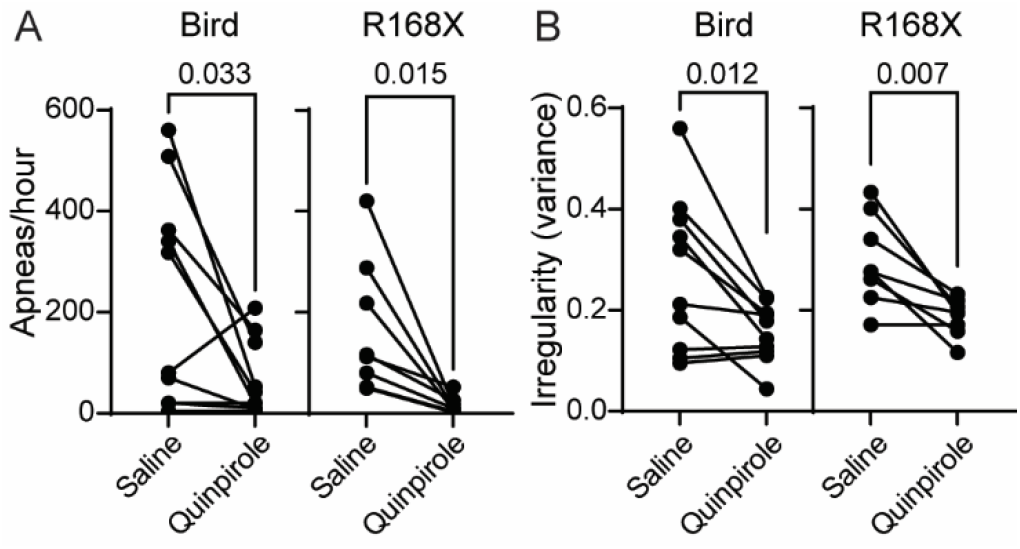
Quinpirole reduced apneas and irregularity score. Whole-body plethysmography data from Mecp2^Bird/+^ or Mecp2^R168X/+^ mice were analyzed 30 – 60 min after saline or quinpirole (1 mg/kg). Solid black circles and connecting lines represent means from individual animals. p values are from paired t-tests. p< 0.05 are considered significant.

**Figure 6.**
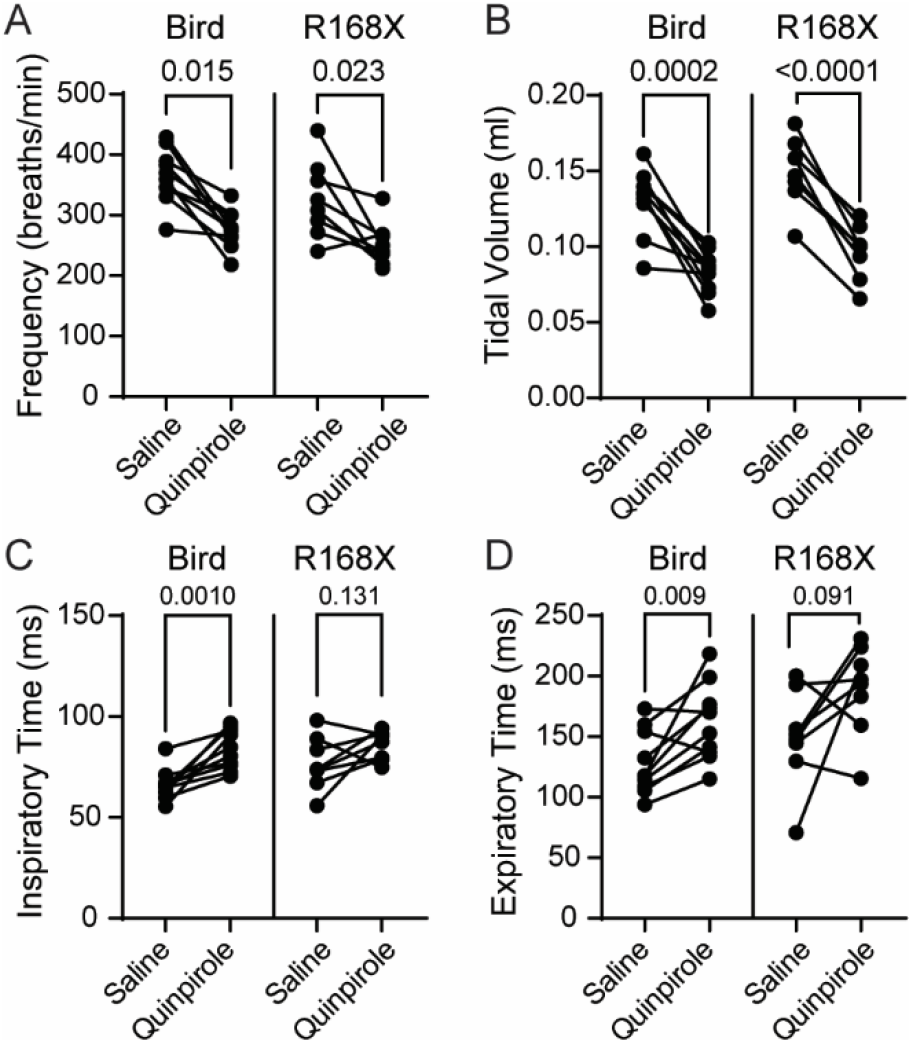
Whole-body plethysmography data from Mecp2^Bird/+^ or Mecp2^R168X/+^ mice. Respiratory parameters (frequency, estimated tidal volume, inspiratory time and expiratory time) were analyzed 30 – 60 min after saline or quinpirole (1 mg/kg). Solid black circles and connecting lines represent means from individual animals. p values are from paired t-tests. p< 0.05 are considered significant.

### Hypercapnic ventilatory responses

Mecp2 deficient mice have an impaired hypercapnic ventilatory response (Zhang *et al*., 2011; Toward *et al*., 2013; Bissonnette *et al*., 2014; Garg *et al*., 2015). We tested the hypercapnic (7% CO_2_) ventilatory response in Mecp2^Bird/+^ and Mecp2^R168X/+^ mice following treatment with saline, PAOPA (1 mg/kg and 10 mg/kg) and quinpirole (1 mg/kg). There were no significant differences in the hypercapnia-induced increase in frequency, tidal volume or minute ventilation between saline and PAOPA-treated mice (Figure 7 A-C). Quinpirole augmented hypercapnia-induced increase in minute ventilation in both Mecp2^Bird/+^ and Mecp2^R168X/+^ mice (Figure 7D). In Mecp2^R168X/+^ mice, this was due to an increase in hypercapnia-induced frequency and tidal volume (Figure 7 E,F). In Mecp2^Bird/+^ mice, there was a significant increase in hypercapnia-induced frequency, but non-significant change in tidal volume (Figure 7 E,F).

**Figure 7.**
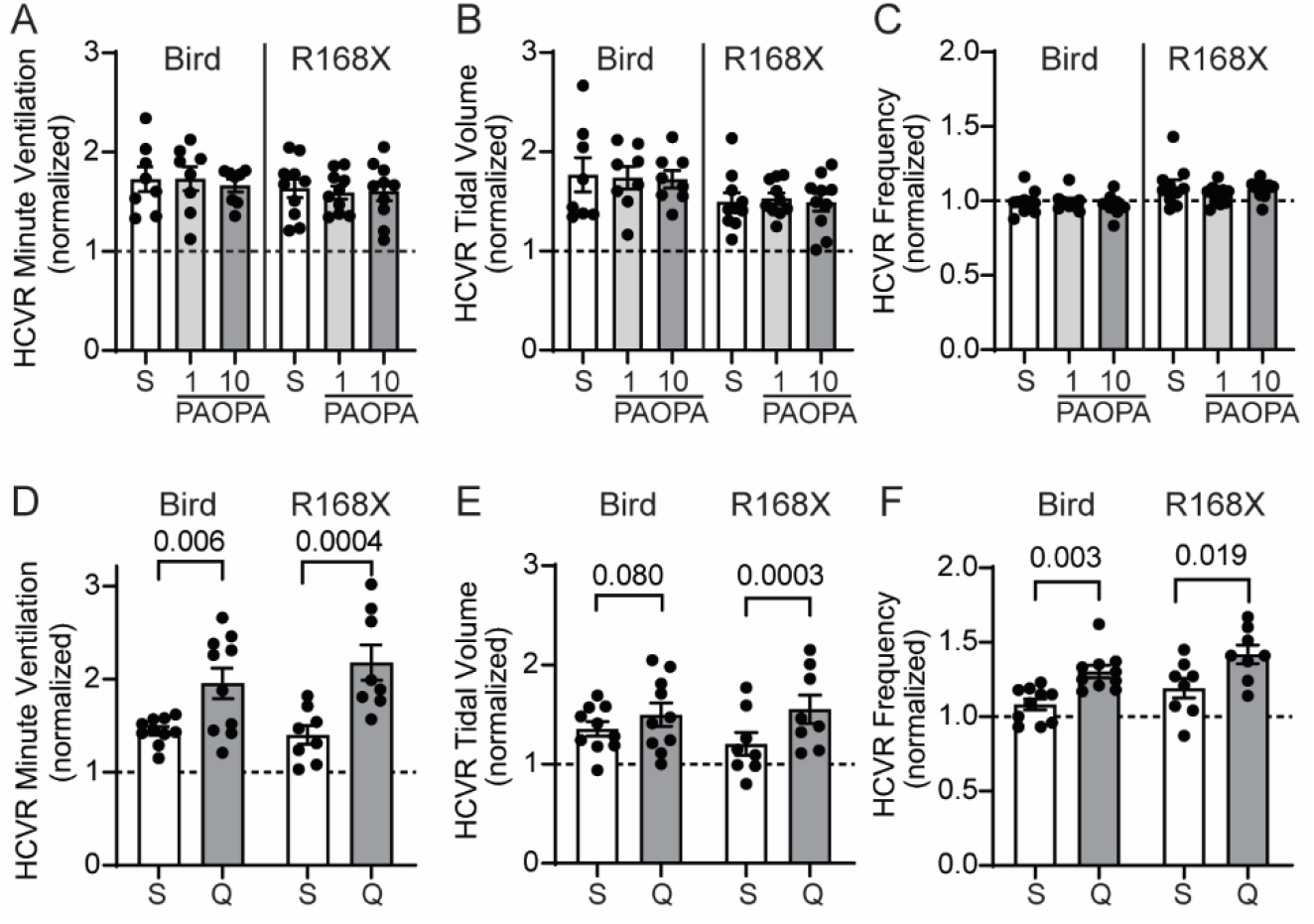
Hypercapnic ventilatory responses (HCVR) from Mecp2^Bird/+^ or Mecp2^R168X/+^ mice. A-C, mice were treated with saline (S) or PAOPA (1 or 10 mg/kg). D-F, mice were treated with saline (S) or quinpirole (Q, 1 mg/kg). Hypercapnia (7% CO2)-induced changes in minute ventilation (A,D), estimated tidal volume (B,E) and frequency (C,F) were normalized to baseline (standard air) levels (baseline = 1, indicated by dashed line). Bars and error represent group mean ± SEM. Solid circles represent individual animals. A-C, no significant differences between saline and PAOPA (p > 0.05 for all comparisons by two-way RM ANOVA and Tukey post-test. D-F, p values from paired t-test; p < 0.05 is considered significant.

### Evaluation of the KF as a D2 Receptor Agonist Site of Action

Overactivity of Kölliker-Fuse (KF) neurons has been proposed as a mechanism of breathing abnormalities in mouse models of RTT (Abdala *et al*., 2015). We used brain slice electrophysiology to determine if dopamine D2 receptor-mediated activation of GIRK conductance could hyperpolarize KF neurons. Whole-cell voltage-clamp recordings were made from KF neurons contained in brain slices from wild-type mice. Dopamine D2 receptor agonists dopamine (10 µM) or quinpirole (1 – 3 µM) did not cause outward GIRK currents in any of the tested KF neurons (Figure 8; n = 7 neurons from 5 mice). GABA-B receptors also activate GIRK channels to hyperpolarize KF neurons (Varga *et al*., 2020). The KF neurons that were tested did have currents following perfusion of GABA-B agonist, baclofen (30 µM), confirming the presence of GIRK channels. Taken together, these data suggest that KF neurons are not directly hyperpolarized by dopamine D2 receptors.

**Figure 8.**
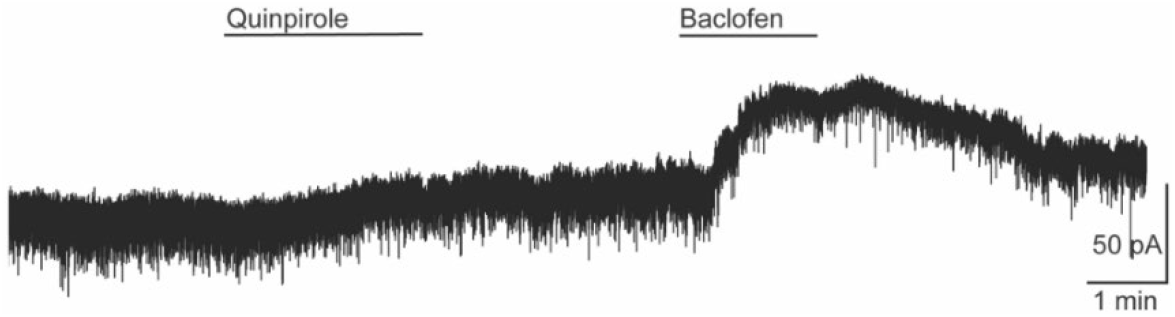
Whole-cell voltage-clamp recording from KF neuron shows absence of quinpirole (1 µM)-mediated current, but presence of baclofen (30 µM)-mediated current in the same neuron.

## Discussion

The present study supports a contribution of dopamine D2 receptors in the previously reported correction of respiratory abnormalities by the 5HT1a and D2-like receptor agonist sarizotan. Although the D2 receptor positive allosteric modulator PAOPA failed to alleviate respiratory perturbations in the Mecp2^R168X/+^ and Mecp2^Bird/+^ mouse models of Rett syndrome, the D2 orthosteric agonist quinpirole dramatically reduced apneas and irregular breathing in mice of both strains. Quinpirole also enhanced the hypercapnic ventilatory response, which is known to be impaired in RTT mice (Zhang et al., 2011; Toward et al., 2013; Bissonnette et al., 2014; Garg et al., 2015).

Quinpirole reduced breathing frequency in both Mecp2^Bird/+^ and Mecp2^R168X/+^ mice. We suspect these respiratory decreases may reflect decreases in locomotor activity and/or body temperature following quinpirole administration (Archer *et al*., 2002; Hofford *et al*., 2012; Luque-Rojas *et al*., 2013). At doses above 0.7 mg/kg, quinpirole reduces locomotion in mice (Archer *et al*., 2002). The present study used a dose of 1 mg/kg, which has been reported to produce an extended period of hypolocomotion in mice (Hofford *et al*., 2012; Luque-Rojas *et al*., 2013). Although our chambers were not equipped to measure locomotor activity, we also observed dramatic reductions in motor behavior following quinpirole administration. Abdala et al. (2013) reported that acute sarizotan treatment produced a depression in locomotor activity and proposed that this was caused by D2 receptor activation in the indirect pathway of the basal ganglia. Pharmacological targeting of indirect pathway neurons containing D2 receptors has been demonstrated to alter locomotor behavior in mice (Muehlmann, et al., 2020). The effect on locomotor activity could also account for some of the reduction in irregularity of breathing, however, it is unlikely to account for the dramatic reduction in apnea frequency, which is a hallmark of the breathing disturbances in RTT mice even in reduced preparations (Abdala *et al*., 2010, 2014). The quinpirole-induced reduction in body temperature and locomotor activity also suggests reductions in metabolic rate, which can stabilize breathing (Gautier, 1996; Haouzi *et al*., 2021). Thus, the stabilization of breathing observed after quinpirole treatment could be secondary to metabolic and locomotor activity changes.

Although PAOPA was ineffective in correcting apnea counts or irregular breathing, our studies of neurophysiology in dopamine neurons of the substantia nigra pars compacta confirm that PAOPA enhances dopamine currents and functions as a positive allosteric modulator at D2 receptors. This physiological finding agrees with cellular and behavioral studies of PAOPA’s selectivity and potency as a D2 positive allosteric modulator (Verma *et al*., 2005; Tan *et al*., 2013). Our observed lack of effects in vivo is also unlikely to be due to insufficient dosing. We used two different doses (1 mg/kg and 10 mg/kg), both of which have been previously reported to have behavioral effects lasting at least an hour following systemic administration in preclinical tests of sensorimotor deficits (Beyaert *et al*., 2013; Daya *et al*., 2018).

Though dopaminergic effects on control of breathing have been reported (Lalley, 2008), the mechanism for dopamine D2 receptor improvements in irregularity, apneas and CO2 sensitivity are unknown. One potential mechanism is inhibition of midbrain dopamine neuron activity through activation of dopamine D2 autoreceptors. Toxin-induced midbrain dopamine neuron depletion reduces breathing frequency and impairs CO2 sensitivity (Tuppy *et al*., 2015; Oliveira *et al*., 2019). We found that quinpirole had similar effects on breathing frequency but enhanced (rather than impaired) CO2 sensitivity, suggesting the effects on CO2 sensitivity are independent of autoreceptor-mediated inhibition of midbrain dopamine neurons. The KF has been localized as a target site in the correction of breathing abnormalities in mouse models of RTT as KF neurons have been shown to be overactive in Mecp2-deficient mice and may contribute to breath holding in RTT (Abdala *et al*., 2014, 2015). However, we found that KF neurons are not directly hyperpolarized by dopamine D2 receptors, implying a lack of postsynaptic D2 receptors in the KF. This does not exclude the possibility of presynaptic D2 receptors in the KF. For example, presynaptic D2 receptors suppress excitatory input onto nearby parabrachial neurons (Chen *et al*., 1999). In addition, D2 receptors are known to suppress the activity of expiratory neurons in the caudal ventral respiratory group (Lalley, 2009).

We observed several MeCP2 mutant mice lacking respiratory symptoms in both models. Genotyping confirmed that these mice were heterozygous for the mutations causing Mecp2 deficiency. Additionally, all respiratory-asymptomatic mice tested displayed hindlimb clasping and obesity – traits that are characteristic of Mecp2-deficiency in mice (Garg *et al*., 2013). Interestingly, body weight was negatively correlated with respiratory symptoms (heavier body weights had less apneas and less irregular breathing). The respiratory asymptomatic mice were all obese. An association between higher body weight and milder symptoms has been reported in RTT patients as well (Renieri *et al*., 2009). It has previously been reported that RTT patients have improvements in abnormal breathing patterns with age (Julu *et al*., 2001; Mackay *et al*., 2017). A similar observation was made by Bissonnette et al. (2014) in the Mecp2^T1158A/+^ mouse model of RTT. The asymptomatic mice in this study were all older than 8 months. Therefore, it is possible that these respiratory-asymptomatic mice underwent similar age-related improvements and were tested too late to observe respiratory defects apparent at earlier points in the lifespan. However, when considered across the entire age range used in the study (5-10 months), age was not correlated with apnea number or irregularity scores, suggesting any age-related improvements were not gradually progressive. The inherent variability of sex-linked disorders due to x-chromosome inactivation may also be responsible for the different phenotypes observed.

In terms of clinical relevance, these data suggest that positive allosteric modulation, at least by PAOPA, may be ineffective in the treatment of respiratory disturbances in RTT. Although orthosteric agonism of the D2 receptor provides a potential therapeutic target for these abnormalities, orthosteric agonists tend to be less selective than their allosteric counterparts, potentially resulting in higher risks of toxicity and incidence of adverse effects (Christopoulos, 2002; Wootten *et al*., 2013; Wold *et al*., 2019). In MPTP-induced monkey models of Parkinson’s disease, quinpirole (and all other D2 agonists tested) alleviated Parkinsonian symptoms, but also reproduced levodopa-induced dyskinesia to a similar degree (Gomez-Mancilla & Bédard, 1991; Pearce *et al*., 1995). Both low and high doses of quinpirole have also been demonstrated to impair reversal learning in marmosets, while intermediate doses improved reversal learning (Horst *et al*., 2019). Chronic quinpirole treatment triggers D2-receptor supersensitization in rats, and has been demonstrated to alter pain thresholds, cause memory deficits, and enhance quinpirole-induced yawning (Kostrzewa, 2017). The potential for side effects is not limited to quinpirole. Dopamine D2-type non-ergoline agonists ropinirole and pramipexole that are in clinical use for Parkinson’s disease also exhibit dopaminergic side effects, including nausea, somnolence and dyskinesias (BROOKS, 2000). These adverse effects present potential barriers to the use of orthosteric D2 receptor agonists in Rett Syndrome.

## Additional Information

### Author contributions

Experiments were performed in the laboratory of Dr. Erica Levitt at the University of Florida. ESL and JMB conceptualized and designed the work. SNB, BTR, DMB, JRW, JTB and ESL performed experiments and data analysis. SNB and ESL drafted the manuscript. All authors critically revised the manuscript, approved the final version of the manuscript, and agree to be accountable for all aspects of the work in ensuring that questions related to the accuracy or integrity of any part of the work are appropriately investigated and resolved. All persons designated as authors qualify for authorship and all those who qualify for authorship are listed.

## Acknowledgements

We would like to thank Natalie Johnson and Jonathan Novak for technical assistance.

